# Polygenic hazard score: an enrichment marker for Alzheimer’s associated amyloid and tau deposition

**DOI:** 10.1101/165373

**Authors:** Chin Hong Tan, Chun Chieh Fan, Elizabeth C. Mormino, Leo P. Sugrue, Iris J. Broce, Christopher P. Hess, William P. Dillion, Luke W. Bonham, Jennifer S. Yokoyama, Celeste M. Karch, James B. Brewer, Gil D. Rabinovici, Bruce L. Miller, Gerald D. Schellenberg, Karolina Kauppi, Howard A. Feldman, Dominic Holland, Linda K. McEvoy, Bradley T. Hyman, Ole A. Andreassen, Anders M. Dale, Rahul S. Desikan, for the Alzheimer’s Disease Neuroimaging Initiative

**Affiliations:** Neuroradiology Section, Department of Radiology and Biomedical Imaging, University of California, San Francisco, San Francisco, California, United States of America; Department of Cognitive Science, University of California, San Diego, La Jolla, California, United States of America; Department of Neurology & Neurological Sciences, Stanford University, Stanford, California, United States of America; Department of Neurology, University of California, San Francisco, San Francisco, California, United States of America; Department of Psychiatry, Washington University in St. Louis, St. Louis, Missouri, United States of America; Department of Neurosciences, University of California, San Diego, La Jolla, California, United States of America; Department of Radiology, University of California, San Diego, La Jolla, California, United States of America; Shiley-Marcos Alzheimer’s Disease Research Center, University of California, San Diego, La Jolla, California, United States of America; Department of Pathology and Laboratory Medicine, University of Pennsylvania, Philadelphia, Pennsylvania, United States of America; Department of Neurology, Massachusetts General Hospital, Boston, Massachusetts, United States of America; NORMENT Institute of Clinical Medicine, University of Oslo and Division of Mental Health and Addiction, Oslo University Hospital, Oslo, Norway

## Abstract

**Background:** There is an urgent need for the early identification of nondemented individuals at the highest risk of progressing to Alzheimer’s disease (AD) dementia for early therapeutic interventions. Our goal was to evaluate whether a recently validated polygenic hazard score (PHS) can be integrated with known *in vivo* CSF or PET biomarkers of amyloid or tau pathology to prospectively predict cognitive decline and clinical progression to AD dementia in nondemented older individuals.

**Methods:** We evaluated 347 cognitive normal (CN) and 599 mild cognitively impaired (MCI) individuals. We first investigated whether PHS can predict CSF or PET amyloid and tau deposition. We evaluated differences in positive and negative predictive values of biomarker status, as a function of PHS risk. Next, we used linear mixed-effects (LME) to examine if PHS and biomarker status in conjunction, best predict longitudinal cognitive and clinical progression. Lastly, we used survival analysis to investigate whether a combination of PHS and biomarker positivity predicts progression to AD dementia better than using PHS or biomarker positivity alone.

**Findings:** In CN and MCI individuals, we found that amyloid and total tau positivity systematically varies as a function of PHS. For individuals in greater than the 50^th^ percentile PHS, the positive predictive value for amyloid approached 100%. Similarly, for individuals in less than the 25^th^ percentile PHS, the negative predictive value for total tau approached 85%. Beyond *APOE*, high PHS individuals with amyloid and tau pathology showed the fastest rate of longitudinal cognitive decline and time to AD dementia progression. Among the CN subgroup, we similarly found that PHS was strongly associated with amyloid positivity and the combination of PHS and biomarker status significantly predicted longitudinal clinical progression.

**Interpretation:** Among asymptomatic and mildly symptomatic older individuals, PHS considerably improves the predictive value of CSF or PET amyloid and tau biomarkers. Beyond *APOE*, PHS may be useful for risk stratification and cohort enrichment for MCI and preclinical AD therapeutic trials.

## INTRODUCTION

Accumulating genetic, molecular, biomarker and clinical evidence indicates that the pathobiological changes underlying late-onset Alzheimer’s disease (AD) occur 20-30 years before the onset of clinical symptoms^1,2^. AD associated pathology may follow a temporal sequence whereby β-amyloid dysmetabolism (i.e. assessed *in vivo* as reductions in CSF levels of Aβ_1-42_ or increase in PET ^18^F-AV-45) precedes tau accumulation (i.e. assessed *in vivo* as elevations in CSF total tau) and neurodegeneration^3^. Given the large societal and clinical impact associated with AD dementia, there is an urgent need to identify and therapeutically target nondemented older individuals with amyloid or tau pathology who may be at greatest risk of progressing to dementia.

A large body of work has shown that genetic risk factors such as the ε4 allele of *apolipoprotein E* (*APOE*) modulate amyloid pathology and shift clinical AD dementia onset to an earlier age^4^. Beyond *APOE* ε4, numerous single nucleotide polymorphisms (SNPs) have now been shown to be associated with small increases in AD dementia risk^5^. Based on a combination of *APOE* and 31 other genetic variants, we have developed and validated a ‘polygenic hazard score’ (PHS) for quantifying AD dementia age of onset.^6^ Importantly, PHS was associated with *in vivo* biomarkers of AD pathology such as reduced CSF Aβ_1-42_ (indicating elevated intracranial amyloid plaques) and elevated CSF total tau (indicating elevated intracranial neurofibrillary tangles). Using a large prospective clinical cohort, we have recently shown that PHS predicts time to AD dementia and longitudinal, multi-domain cognitive decline in cognitively normal (CN) individuals and in patients with mild cognitive impairment (MCI)^7^.

However, the value of combining PHS with *in vivo* biomarkers of AD pathology to predict cognitive and clinical progression among nondemented older individuals remains unknown. Here, among MCI and CN individuals, we evaluated whether PHS can be useful as a marker for enriching and stratifying Alzheimer’s associated amyloid and tau pathology.

## METHODS

### Participants and clinical characterization

We evaluated individuals with longitudinal genetic, clinical, neuropsychological, PET and CSF measurements from the Alzheimer’s Disease Neuroimaging Initiative 1, GO, and 2 (ADNI1, ADNI-GO, and ADNI2). Data used in the preparation of this article were obtained from the Alzheimer’s Disease Neuroimaging Initiative (ADNI) database (adni.loni.usc.edu). The ADNI was launched in 2003 as a public-private partnership, led by Principal Investigator Michael W. Weiner, MD. The primary goal of ADNI has been to test whether serial magnetic resonance imaging (MRI), positron emission tomography (PET), other biological markers, and clinical and neuropsychological assessment can be combined to measure the progression of mild cognitive impairment (MCI) and early Alzheimer’s disease (AD). We restricted analyses to CN individuals (*n* = 347) and patients diagnosed with MCI, (*n* = 599) who had both genetics and CSF or PET biomarkers (CSF Aβ_1-42_, CSF total tau, or PET ^18^F-AV45) data. We used previously established thresholds of <192 pg/ml, >23 pg/ml^8^ and >1.1^9^ to indicate ‘positivity’ for CSF Aβ_1-42_, CSF total tau and PET ^18^F-AV-45, respectively. We classified individuals as amyloid ‘positive’ if they reached either CSF Aβ_1-42_ or PET ^18^F-AV45 threshold for positivity, given that PET and CSF amyloid biomarkers provide highly correlated measurements of intracranial amyloid deposition^10^. Cohort demographics are summarized in Table 1.

**Table 1.**
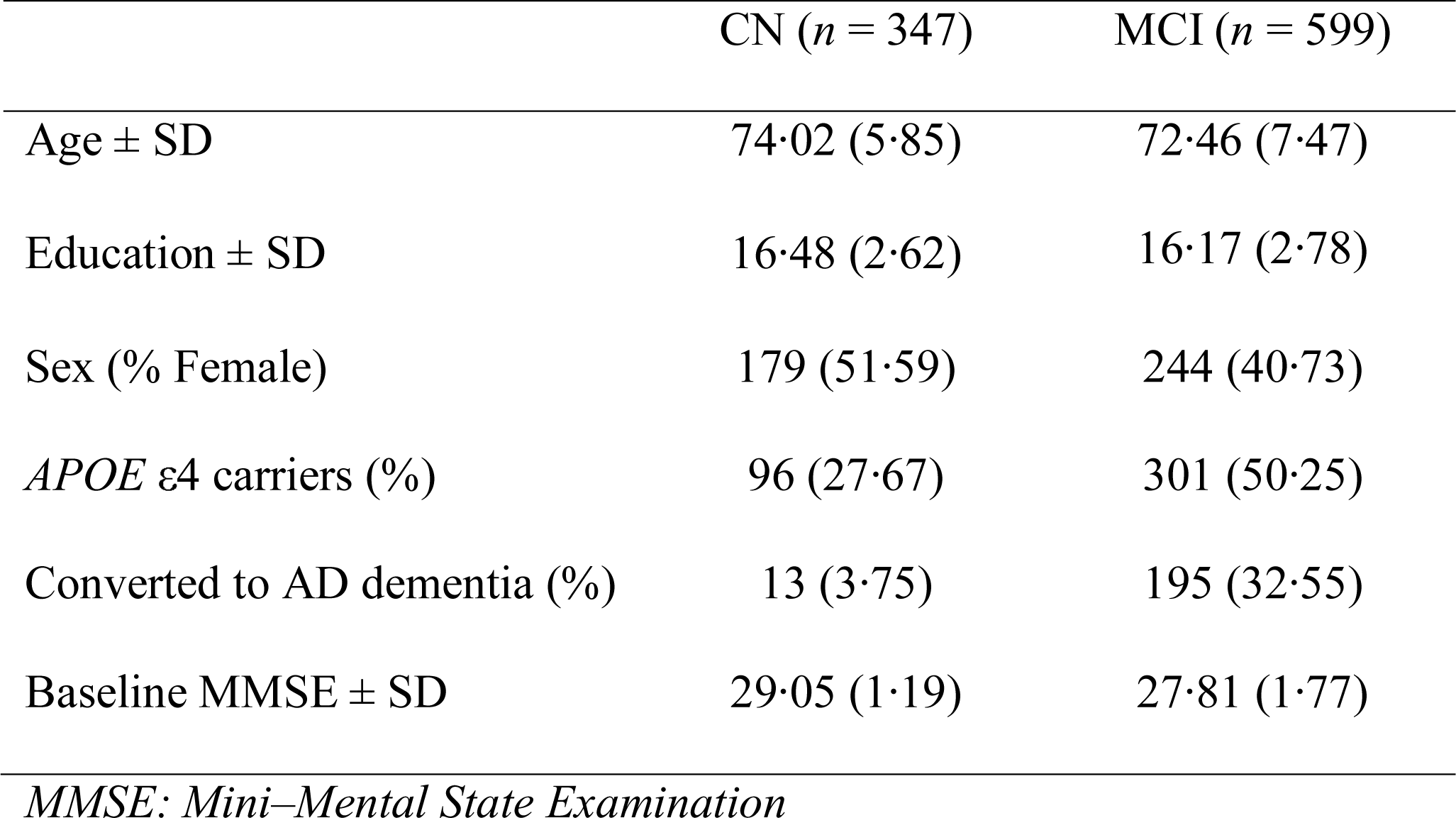
Cohort demographics

### Polygenic hazard score (PHS)

For each CN and MCI participant in this study, we calculated their individual PHS, as previously described^6^. In brief, AD associated SNPs (at *p* < 10^-5^) were first identified using genotype data from 17,008 AD cases and 37,154 controls from Stage 1 of the International Genomics of Alzheimer’s Disease Project. Next, these AD associated SNPs (final total of 31 SNPs) were selected based on stepwise procedures of Cox proportional hazards models using genotype data from 6,409 AD patients and 9,386 older controls from Phase 1 of the Alzheimer’s Disease Genetics Consortium (ADGC Phase 1), providing a polygenic hazard score (PHS) for each participant. Finally, by combining US population based incidence rates, and genotype-derived PHS for each individual, estimates of instantaneous risk for developing AD, based on genotype and age, were derived. In this study, the PHS computed for every CN and MCI participant represents the vector product of an individual’s genotype for the 31 SNPs and the corresponding parameter estimates from the ADGC Phase 1 Cox proportional hazard model.

### Statistical analysis

Using logistic regression, we first evaluated the relationship between PHS and baseline amyloid and total tau positivity (binarized as positive or negative) in CN and MCI individuals. In these analyses, we controlled for age at baseline, sex, education, and *APOE* ε4 status (binarized as having at least one copy of the ε4 allele versus none). We further ascertained the ‘enrichment’ in amyloid and tau positive predictive value (PPV), and negative predictive value (NPV) for clinical progression to AD dementia, as a function of PHS percentiles.

Next, we used linear mixed-effects (LME) models in CN and MCI individuals to investigate whether a statistical interaction between PHS and amyloid or total tau status significantly predicted longitudinal cognitive decline and clinical progression (mean follow-up time = 2.36 years, SD = 1.99 years). We conducted LME models separately for amyloid and total tau. We defined cognitive decline using change scores in 2 domains, namely executive function and memory, based on composite scores developed using the ADNI neuropsychological battery and validated using confirmatory factor analysis^11^. We defined clinical progression using change scores in Cognitive Dementia Rating - Sum of Boxes (CDR-SB). We examined the main and interactive effects of PHS and biomarker positivity on cognitive decline and clinical progression rate while controlling for age, sex, education, *APOE* ε4 status, using the following LME model:

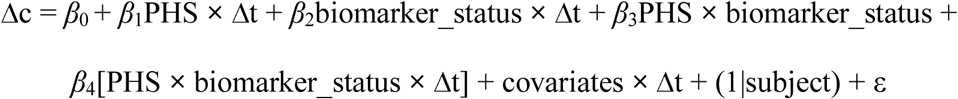

Here, Δc = cognitive decline (executive function or memory) or clinical progression (CDR-SB) rate, Δt = change in time from baseline visit (years), biomarker_status = positive or negative for amyloid or total tau, and (1|subject) specifies the random intercept. We centered and scaled the continuous predictors (PHS and time) and covariates (age at baseline and education) prior to analysis. We were specifically interested in PHS **×** biomarker_status **×** Δt, whereby a significant interaction indicates differences in rates of decline, as a function of differences in PHS and biomarker status. We then examined the simple main effects by comparing slopes of cognitive decline and clinical progression over time for individuals who were biomarker positive or negative, and with either high (∼84 percentile) or low PHS (∼16 percentile). We defined high and low PHS by 1 standard deviation above or below the mean of PHS respectively^7^. To assess the added utility of PHS in predicting cognitive and clinical decline, we further compared the LME models with reduced LME models without PHS using likelihood ratio tests.

Using a survival analysis framework, we next investigated the value of combining PHS with amyloid and total tau biomarkers status to predict time to AD dementia progression. Specifically, we examined the effects of 1) PHS, 2) PHS in individuals who were amyloid positive, and 3) PHS in individuals who were amyloid and total tau positive, on time to AD dementia progression using a Cox proportional hazards model. Age of AD dementia onset was used as the time to event. ‘Ties’ were resolved using the Breslow method. We co-varied for the effects of sex, education, *APOE* ε4 status, age at baseline, and also age at baseline stratified into quintiles to adjust for violations of Cox proportional hazards assumptions by baseline age. We restricted survival analyses, which involves AD dementia censoring, as well as the PPV/NPV analyses (see above) to the combined CN and MCI groups only as the CN individuals had low conversion rates during the observation period (< 5%).

Finally, building on our prior work^6^, we assessed whether amyloid and tau status could inform PHS-predicted annualized incidence rate of AD age of onset. We examined the influence of amyloid status, total tau status, and both in combination on the PHS-derived annualized incidence rate of AD dementia age of onset, based on previously established AD incidence estimates from the United States population^12^.

## RESULTS

### PHS enriches AD predictive value of amyloid and tau deposition

Within the combined MCI and CN cohort, we found that PHS predicted amyloid (Odds ratio (OR) = 2·31, 95% Confidence Interval (CI) = 1·55 – 3·49, *p* = 5·50×10^-5^) and total tau (OR =1·87, 95% CI = 1·31 - 2·68, *p* = 6·53×10^-4^) positivity. As illustrated in Figure 1, we found that the proportion of individuals who were amyloid or total tau positive increased systematically as a function of higher PHS. For example, approximately 60% of individuals in the 75^th^ PHS percentile would be classified as amyloid positive whereas less than 40% of individuals in the 25^th^ PHS percentile would be amyloid positive. In subgroup analyses involving only CN individuals, PHS predicted amyloid (OR = 2·44, 95% CI = 1·13 – 5·55, *p* = 0·028) but not total tau (OR =1·53, 95% CI = 0·71 – 3·35, *p* = 0·28) positivity. Within MCI individuals only, PHS predicted amyloid (OR = 1·96, 95% CI = 1·17 – 3·33, *p* = 1·12×10^-2^) and total tau (OR =2·24, 95% CI = 1·12 - 4·59, *p* = 2·46×10^-2^) positivity.

**Figure 1.**
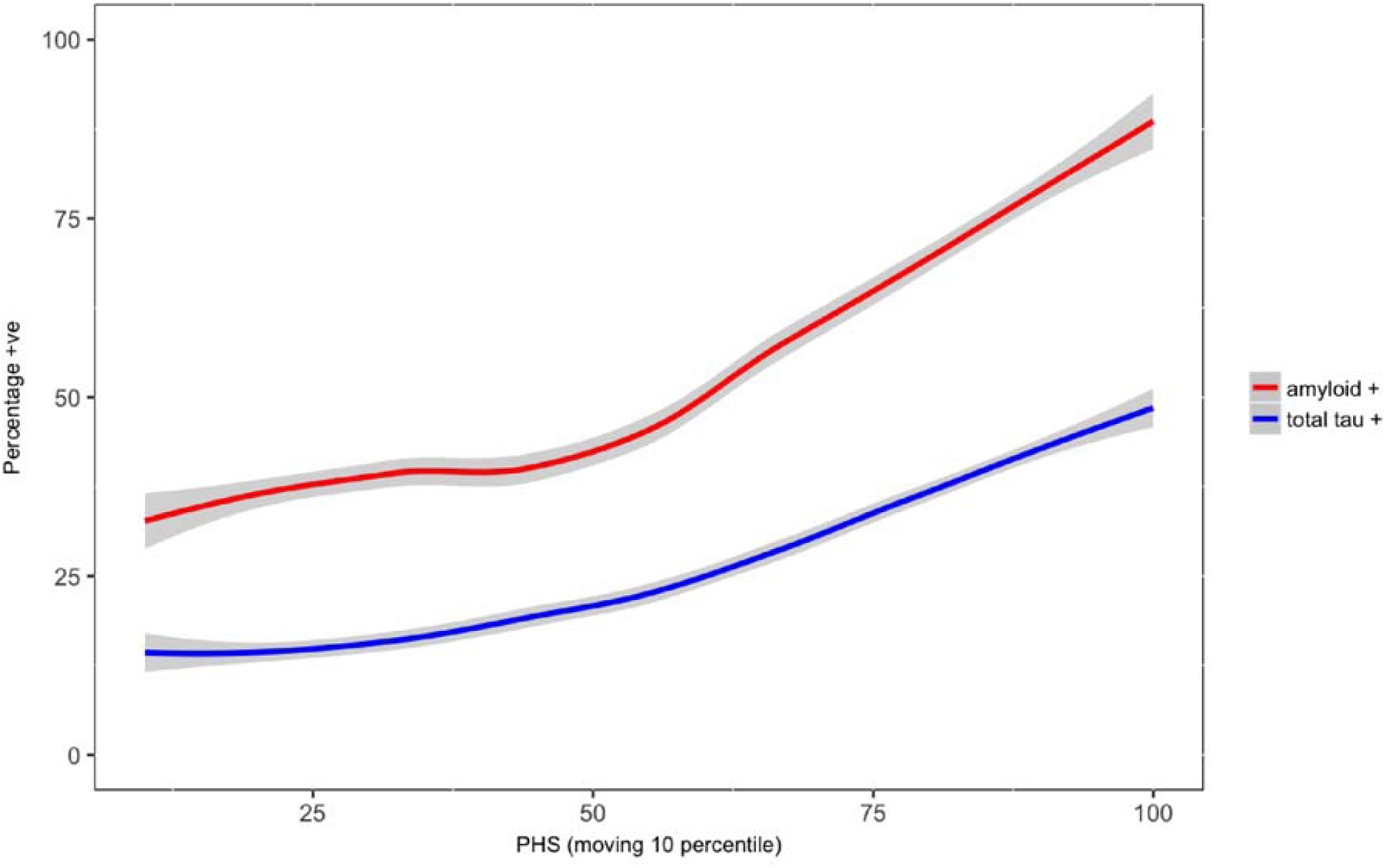
Increase in proportion of nondemented individuals who tested positive on amyloid (red) and total tau (blue) as a function of higher polygenic hazard score (moving 10 percentiles, 1% increment per step).

Similarly, within the combined MCI and CN cohort, we found that PPV of amyloid and total tau increased systematically as a function of higher PHS percentiles (Figure 2a). For instance, the PPV for amyloid positivity approaches 100% amongst individuals with ≥50th percentile PHS but for individuals in ≤ 25^th^ percentile PHS, amyloid PPV is approximately 75%. Based on a 1000 bootstrap of 50 random samples, PPV for all individuals was higher for amyloid compared to total tau (Welch t-test, *t* (1339·1) = 137·92, *p* <2×10^-16^). In contrast, we found that NPV was highest in individuals with low PHS percentiles, especially for total tau (Figure 2b). For amyloid and total tau, we note that the maximum absolute value PPV (approximately 98%) as a function of PHS was higher than the maximum absolute NPV (approximately 85%) as a function of PHS.

**Figure 2.**
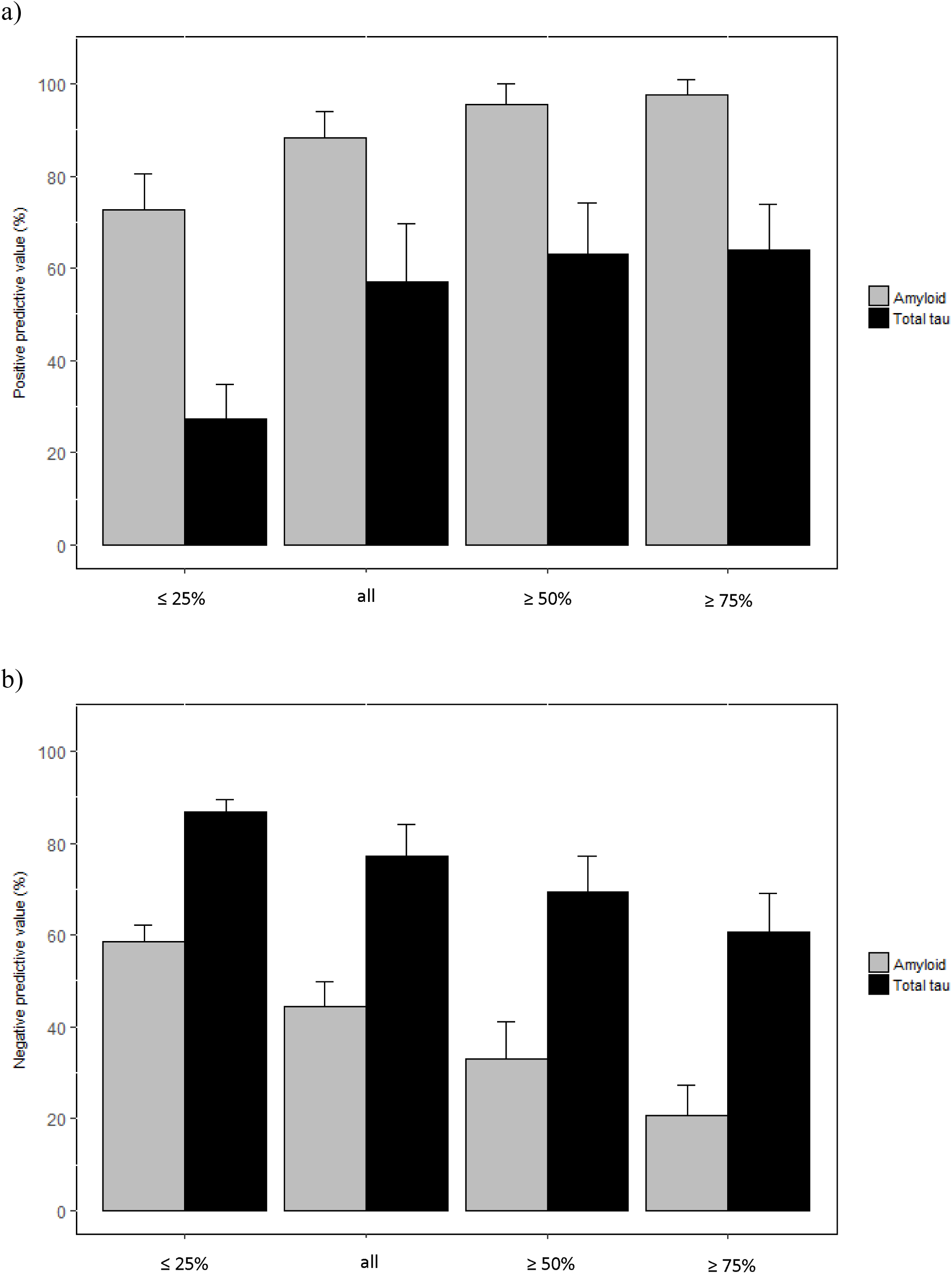
a) Positive predictive value (PPV) and b) negative predictive value (NPV) of amyloid and total tau with subsequent progression to AD dementia, based on stratification of polygenic hazard score (PHS) into different percentile bins (≤ 25%, all individuals, ≥ 50% and ≥ 75%).Error bars are 1000 bootstrap estimate of the standard deviation of 50 random samples in each PHS percentile bins.

### PHS enriches amyloid and tau associated prediction of clinical and cognitive decline

Using LME analyses within the combined MCI and CN cohort, we investigated the role of PHS in conjunction with amyloid and total tau positivity in predicting cognitive decline (executive function and memory) and clinical progression (i.e. increase in CDR-SB), and found that the 3-way interactions (PHS **×** biomarker_status **×** Δt) were statistically significant for amyloid in executive function (β = -0·14, SE = 0·02, *p* = 1·01×10^-13^), memory (β = -0·05, SE = 0·02, *p* = 1·87×10^-3^), CDR-SB: β = 0·55, SE = 0·06, *p* < 2×10^-16^), and for total tau in executive function (β = -0·10, SE = 0·02, *p* = 1·39×10^-9^), memory (β = -0·05, SE = 0.02, *p* =1·24×10^-3^), and CDR-SB (β = 0·40, SE = 0·06, *p* = 2·35×10^-12^). We conducted simple slopes analyses and found that individuals who had high PHS (∼84 percentile) and tested positive for amyloid experienced the fastest rate of cognitive decline (executive function: β = -0·43, SE = 0·04, *p* < 2×10^-16^; memory: β = -0·37, SE = 0·03, *p* <2×10^-16^) and clinical progression (CDR-SB: β = 1·72, SE = 0·12, *p* <2×10^-16^). Similarly, individuals with high PHS and positive for total tau also experienced the fastest rate cognitive and clinical decline (executive function: β = -0·46, SE = 0·04, *p* < 2×10^-16^; memory: β = -0·38, SE = 0·03, *p* < 2×10^-16^; CDR-SB: β = 1·86, SE = 0·13, *p* < 2×10^-16^ (Supplementary Table 1, Figure 3).

**Figure 3.**
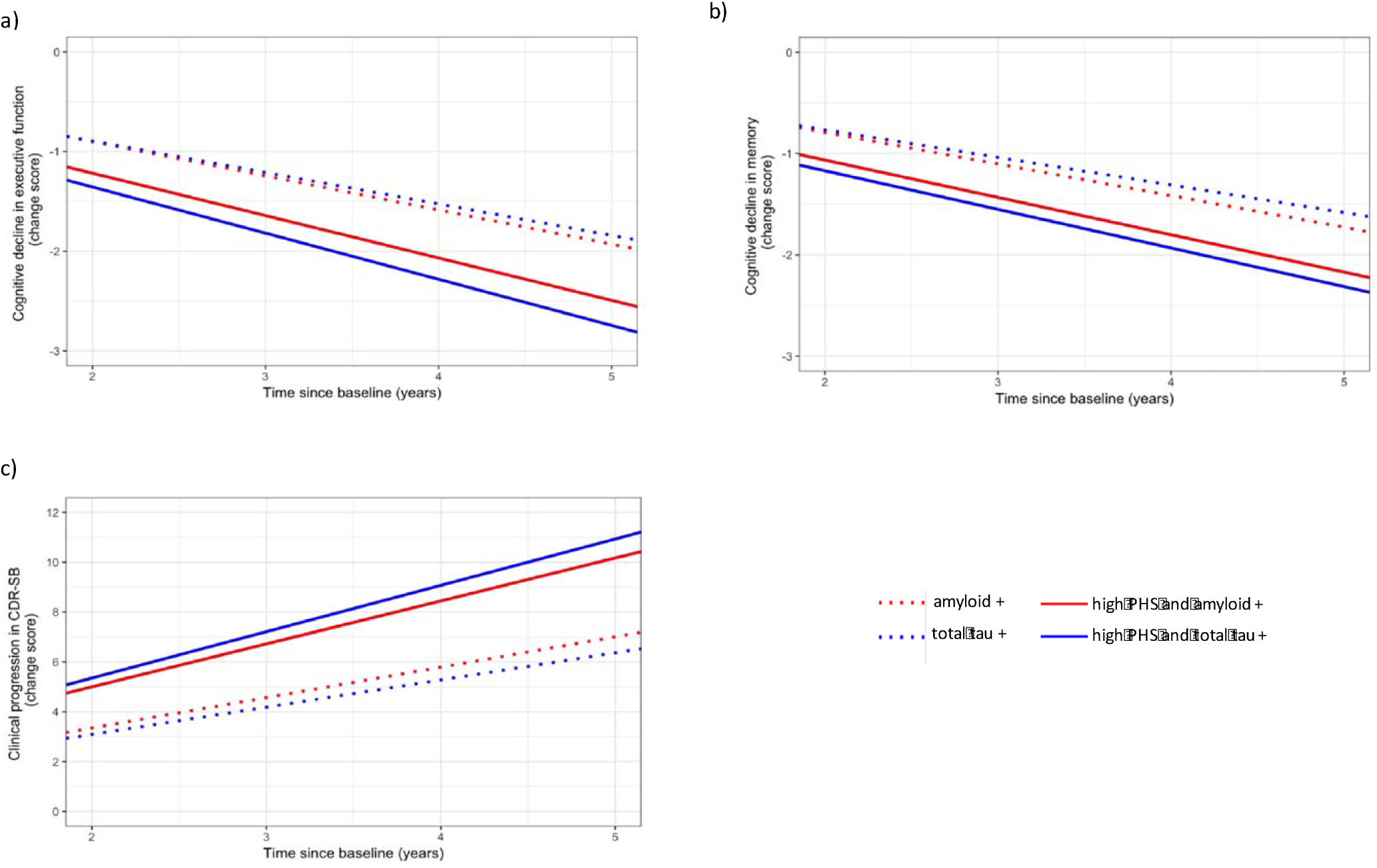
Differences in rates of cognitive decline in a) executive function, b) memory, and clinical progression in c) cognitive dementia rating – sum of boxes (CDR-SB) over time for high polygenic hazard score (PHS) individuals who tested positive for amyloid (solid red line) or total tau (solid blue line) in full PHS linear mixed-effects (LME) models, compared to individuals who tested positive for amyloid (dotted red line) or total tau (dotted blue line) in reduced non-PHS LME models (see text for model details).

In addition, using likelihood ratio tests, we found that these full LME models resulted in a better fit than a reduced, non-PHS model with only amyloid status (executive function: χ^2^(4) = 76·3, *p* = 1·07**×**10^-15^; memory: χ^2^(4) = 34·3, *p* = 6·45**×**10^-7^; CDR-SB: χ^2^(1) = 143·71, *p* < 2**×**10^-16^) or total tau status (executive function: χ^2^(4) = 56·3, *p* = 1·76**×**10^-11^; memory: χ^2^(4) = 42·2, *p* = 1·54**×**10^-8^; CDR-SB: χ^2^(1) = 113·11, *p* < 2**×**10^-16^). Specifically, amyloid or total tau positive individuals with high PHS showed greater cognitive decline and faster clinical progression than individuals who were amyloid or total tau positive, regardless of PHS (Figure 3). These findings demonstrate the added value of using PHS in conjunction with biomarker status to identify individuals who would experience the greatest rate of cognitive decline and clinical progression.

In CN subgroup analyses, the 3-way interaction was only significant for PHS and amyloid status in predicting change in CDR-SB (β = 0·16, SE = 0·07, *p* = 1·88×10^-2^), but not executive function (β = -0·02, SE = 0·03, *p* = 0.42) and memory (β = 0·04, SE = 0.03, *p* = 0·11). In simple main effects analysis for CDR-SB, high PHS and amyloid positive individuals also showed the greatest rate of progression (β = 0·60, SE = 0·13, *p* = 3·47×10^-6^). For PHS with total tau status, the 3-way interaction was significant in predicting changes in executive function (β = -0·11, SE = 0·03, *p* = 5·77×10^-4^) and CDR-SB (β = 0·30, SE = 0·08, *p* = 9·21×10^-5^) but not memory (β = -0·03, SE = 0·03, *p* = 0·36). Similarly, in simple main effect analyses, high PHS and tau positive individuals showed the greatest rate of cognitive and clinical decline (executive function: β = -0·28, SE = 0·06, *p* = 7·26×10^-6^; CDR-SB: β = 0·85, SE = 0·15, *p* = 4·18×10^-8^).

In MCI subgroup analyses, the 3-way interaction was significant for PHS and amyloid status in predicting change in executive function (β = -0·14, SE = 0·02, *p* = 3·47×10^-9^), memory (β = -0·08, SE = 0·02, *p* = 2·22×10^-4^), and CDR-SB (β = 0·49, SE = 0.09, *p* = 2·29×10^-8^). In simple main effect analysis, high PHS and amyloid positive individuals also showed the greatest rate of change for executive function (β = -0·51, SE = 0·04, *p* < 2×10^-16^), memory (β = -0·42, SE = 0·04, *p* < 2×10^-16^) and CDR-SB (β = 2·24, SE = 0·17, *p* < 2×10^-16^). For PHS with total tau status, the 3-way interaction was significant in predicting changes in CDR-SB (β = 0·17, SE = 0·07, *p* = 1·94×10^-2^) but not for executive function (β = -0·03, SE = 0·02, *p* = 0·14) and memory (β = -0·03, SE = 0·02, *p* = 0·11). In simple main effect analyses of the significant interaction, high PHS and tau positive individuals showed the greatest rate of clinical progression (CDR-SB: β = 2·25, SE = 0·17, *p* < 2×10^-16^).

### PHS enriches amyloid + tau prediction of time to AD dementia progression

We used Cox proportional hazards model to investigate clinical progression of nondemented individuals (CN + MCI) to AD dementia, and found that PHS predicted progression (Hazard ratio (HR) = 1·74, 95% CI = 1·30 – 2·33, *p* = 2·26**×**10^-4^). PHS also predicted progression when including amyloid status in the model (HR = 1·49, 95% CI = 1·11 – 2·00, *p* = 8·06**×**10^-3^), and when including both amyloid and total tau status in conjunction (HR = 1·44, 95% CI = 1·07 – 1·94, *p* = 1·67**×**10^-2^)(Figure 4). Comparing goodness of fit using likelihood ratio tests, we found that the reduced models involving only PHS (and covariates) were improved when including amyloid status (χ^2^(1) = 54·1, *p* = 1·94**×**10^-13^), and both amyloid status and total tau status in conjunction (χ^2^(2) = 80·1, *p* < 2**×**10^-16^). The combined model which included PHS, amyloid and total tau status showed the best fit and best predicted time to AD dementia progression than just amyloid and tau status (χ^2^(1) = 5·59, *p* = 1·81**×**10^-2^).

**Figure 4.**
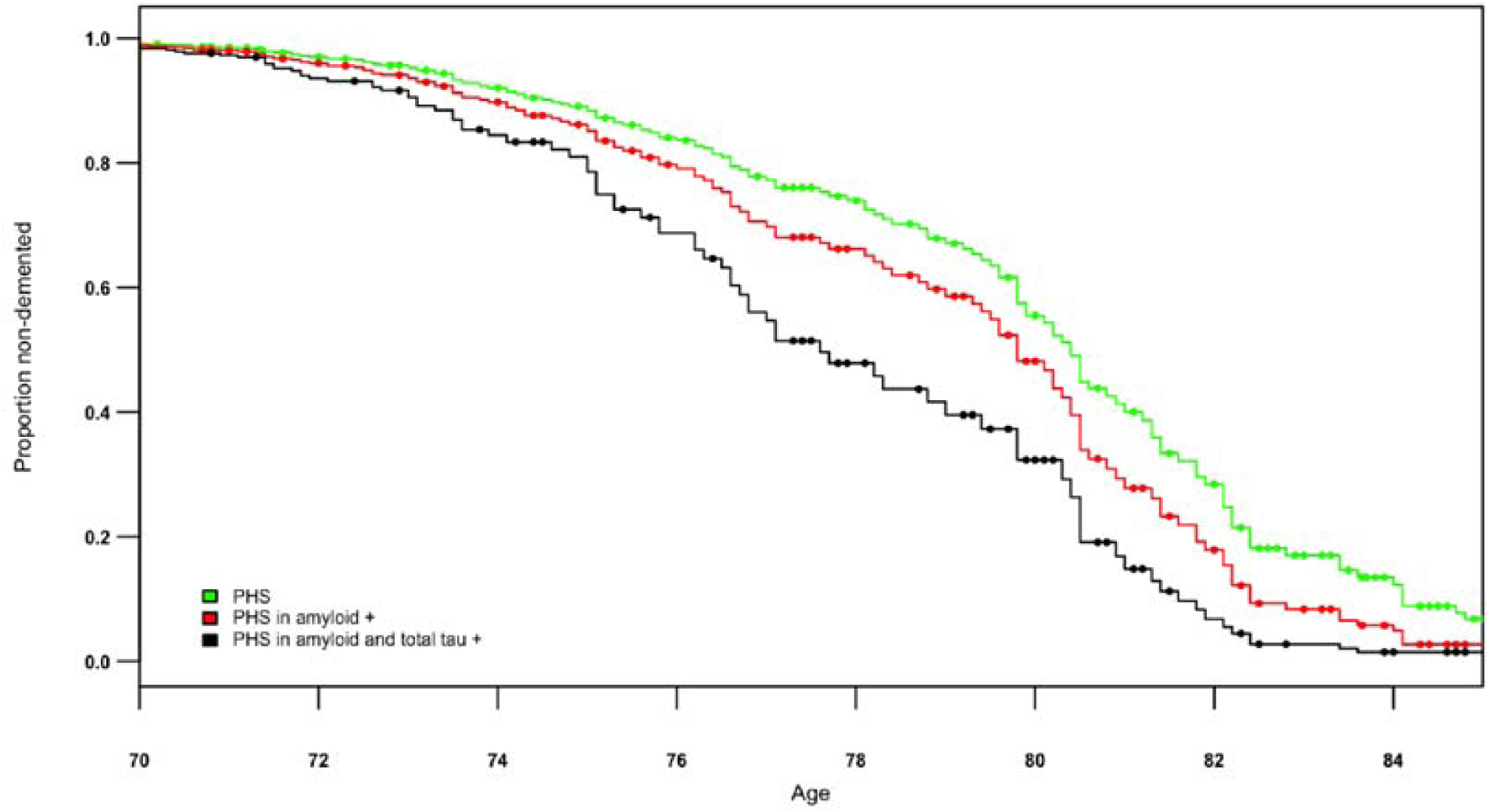
Survivor plot showing greater progression to AD dementia for all individuals as a function of polygenic hazard score (PHS) in individuals who tested positive for both amyloid and total tau (black), compared to subset of individuals who tested positive for amyloid (red), and PHS alone without biomarker information (green).

### Amyloid + tau positive individuals show highest PHS derived annualized AD incidence rates

Finally, we generated population baseline-corrected survival curves stratified by amyloid and total tau positivity status and converted them directly into incidence rates based on PHS^6^. This measure of cumulative incidence rate (CIR) based on age and PHS provides the annualized risk of a nondemented individual for progressing to AD dementia. As illustrated in Figure 5, we found that amyloid and total tau positive individuals showed the highest CIRs compared to other groups, particularly at later ages (over 80). An individual who tested negative for amyloid at baseline would have a CIR of 0.025 at age 70 and 0.37 at age 90. In contrast, an individual who tested positive for both amyloid and total tau at baseline would have a higher CIR of 0.075 at age 70, and 0.73 at age 90. These results indicate that amyloid and total tau positive individuals with high PHS risk will likely experience the highest annualized AD dementia incidence rates.

**Figure 5.**
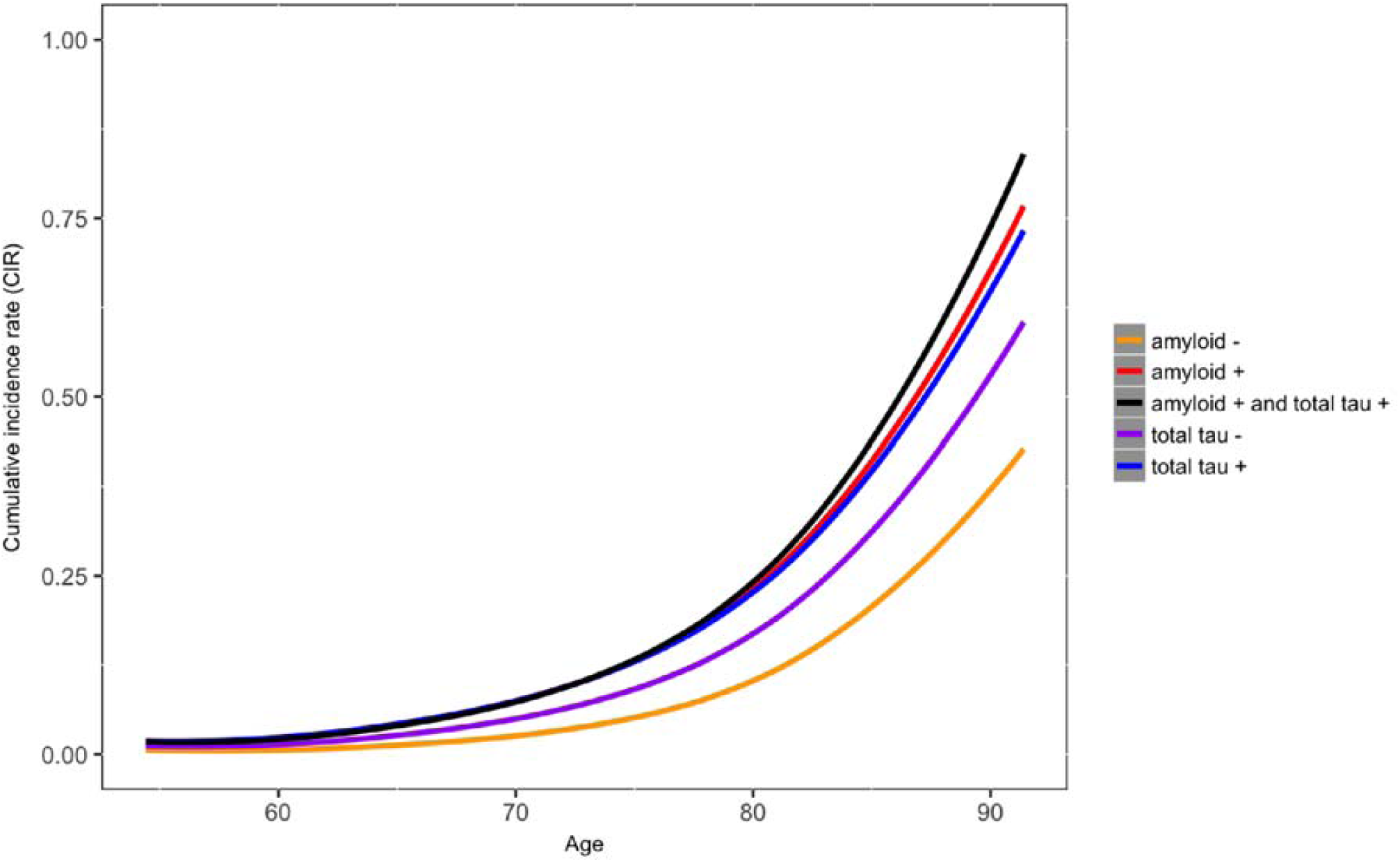
Annualized cumulative incidence rates depicting instantaneous hazard based on an individual’s age and polygenic hazard risk score, stratified based on individuals who were amyloid positive (red), amyloid negative (orange), tau positive (blue), tau negative (purple), and both amyloid and tau positive (black).

## DISCUSSION

In this study, we show that the predictive value of Alzheimer’s associated intracranial amyloid and tau deposition increases systematically as a function of PHS; for MCI and CN individuals in greater than the 50^th^ percentile PHS, the positive predictive value for amyloid approached 100%. Beyond *APOE* ε4, by integrating PHS with *in vivo* biomarkers of AD pathology, we were able to identify nondemented older individuals at highest risk for AD dementia progression and with the steepest longitudinal cognitive decline. Crucially, the combination of PHS, amyloid and total tau best predicted time to AD dementia progression and cognitive decline further indicating the value of stratifying by PHS, beyond amyloid or total tau alone. Collectively, our findings indicate that among nondemented older individuals, PHS can serve as an ‘enrichment’ marker for AD associated intracranial amyloid and tau deposition.

From a clinical trial perspective, PHS may be useful in AD secondary prevention and therapeutic trials. Building on prior work^13,14^, our PPV results indicate that PHS can be used to enrich clinical trial cohorts by identifying those nondemented individuals with amyloid or tau pathology who are at highest risk of progressing to AD dementia and therefore most likely to benefit from therapeutic intervention. As PHS strongly predicted both amyloid and total tau positivity, PHS may also serve as an initial screening tool for *in vivo* AD pathology; for cohort enrichment in trials, it may be helpful to pursue a stratified approach^15,16^ where costlier secondary assessments using CSF or PET biomarker assays are obtained only in those individuals with a high PHS. We note that the PPV was higher for amyloid compared to total tau indicating that PHS provides more enrichment in amyloid compared to tau pathology. In contrast, we found that NPV was highest in individuals with low PHS percentiles, especially for total tau, indicating that individuals who tested negative for total tau and had low PHS were very unlikely to progress to AD dementia subsequently.

From a clinical perspective, PHS may be useful for risk stratification of nondemented older individuals. In our linear mixed effects and survival analyses, we found that PHS considerably improved the ability of amyloid and tau to predict time to AD dementia progression and longitudinal cognitive decline. Importantly, the combination of PHS, amyloid and total tau best predicted clinical and cognitive decline. Together, these findings suggest that a PHS- stratified approach may be clinically useful. Therefore, PHS can be useful as a first step for determining an older individual’s genetic risk for AD dementia. Rather than evaluating all individuals, amyloid and tau assessments may be most helpful only in those individuals with a high PHS. Conversely, among individuals with a low PHS, it may be less effective to pursue additional evaluation with amyloid biomarkers.

Building on prior work evaluating polygenic risk in preclinical AD^17,18,19^ our findings indicate that PHS may be useful for predicting clinical and cognitive decline among asymptomatic older individuals. Among CNs, we found that significant interactions between PHS and amyloid or tau predicted longitudinal change in CDR-SB. Furthermore, PHS predicted amyloid positivity even in CN individuals. Together, these results indicate that the utility of using PHS for cohort enrichment and pre-screening may extend into preclinical AD. In other words, PHS can identify cognitively asymptomatic individuals who are more likely to show AD pathology and who may be at highest risk for clinical progression over time.

In conclusion, we show the utility of integrating PHS with *in vivo* biomarkers of amyloid and tau pathology for cohort enrichment in clinical trials and risk stratification for MCI and preclinical AD. Beyond *APOE*, our findings indicate that stratification by PHS considerably ‘boosts’ the predictive value of amyloid. Among nondemented older individuals of European ancestry, the combination of both PHS and biomarker status best predicts cognitive and clinical decline. Future work should evaluate the value of PHS as an AD associated risk stratication and cohort enrichment marker in diverse, non-Caucasian, non-European populations.

## ACKNOWLEDGEMENTS

We thank the Shiley-Marcos Alzheimer’s Disease Research Center at UCSD, UCSF Memory and Aging Center and UCSF Center for Precision Neuroimaging for continued support. This work was supported by the RSNA Resident/Fellow Award, ASNR Foundation AD Imaging Award, NACC JI award, National Institutes of Health grants (NIH-AG046374, K01AG049152), the Research Council of Norway (#213837, #225989, #223273, #237250/EU JPND), the South East Norway Health Authority (2013-123), Norwegian Health Association, the KG Jebsen Foundation and AG046374 (CMK). Please see Supplemental Acknowledgements for ADNI, NIAGADS and ADGC funding sources.

**Declaration of interests:** JBB served on advisory boards for Elan, Bristol-Myers Squibb, Avanir, Novartis, Genentech, and Eli Lilly and holds stock options in CorTechs Labs, Inc. and Human Longevity, Inc. AMD is a founder of and holds equity in CorTechs Labs, Inc., and serves on its Scientific Advisory Board. He is also a member of the Scientific Advisory Board of Human Longevity, Inc. (HLI), and receives research funding from General Electric Healthcare (GEHC). The terms of these arrangements have been reviewed and approved by the University of California, San Diego in accordance with its conflict of interest policies.

## SUPPLEMENTAL ACKNOWLEDGEMENTS

**ADNI:** Data collection and sharing for this project was funded by the Alzheimer’s Disease Neuroimaging Initiative (ADNI) (National Institutes of Health Grant U01 AG024904) and DOD ADNI (Department of Defense award number W81XWH-12-2-0012). ADNI is funded by the National Institute on Aging, the National Institute of Biomedical Imaging and Bioengineering, and through generous contributions from the following: AbbVie, Alzheimer’s Association; Alzheimer’s Drug Discovery Foundation; Araclon Biotech; BioClinica, Inc.; Biogen; Bristol-Myers Squibb Company; CereSpir, Inc.; Cogstate; Eisai Inc.; Elan Pharmaceuticals, Inc.; Eli Lilly and Company; EuroImmun; F. Hoffmann-La Roche Ltd and its affiliated company Genentech, Inc.; Fujirebio; GE Healthcare; IXICO Ltd.; Janssen Alzheimer Immunotherapy Research & Development, LLC.; Johnson & Johnson Pharmaceutical Research & Development LLC.; Lumosity; Lundbeck; Merck & Co., Inc.; Meso Scale Diagnostics, LLC.; NeuroRx Research; Neurotrack Technologies; Novartis Pharmaceuticals Corporation; Pfizer Inc.; Piramal Imaging; Servier; Takeda Pharmaceutical Company; and Transition Therapeutics. The Canadian Institutes of Health Research is providing funds to support ADNI clinical sites in Canada. Private sector contributions are facilitated by the Foundation for the National Institutes of Health (www.fnih.org). The grantee organization is the Northern California Institute for Research and Education, and the study is coordinated by the Alzheimer’s Therapeutic Research Institute at the University of Southern California. ADNI data are disseminated by the Laboratory for Neuro Imaging at the University of Southern California.

**ADGC:** The National Institutes of Health, National Institute on Aging (NIH-NIA) supported this work through the following grants: ADGC, U01 AG032984, RC2 AG036528; NACC, U01 AG016976; NCRAD, U24 AG021886; NIA LOAD, U24 AG026395, U24 AG026390; Banner Sun Health Research Institute P30 AG019610; Boston University, P30 AG013846, U01 AG10483, R01 CA129769, R01 MH080295, R01 AG017173, R01 AG025259, R01AG33193; Columbia University, P50 AG008702, R37 AG015473; Duke University, P30 AG028377, AG05128; Emory University, AG025688; Group Health Research Institute, UO1 AG06781, UO1 HG004610; Indiana University, P30 AG10133; Johns Hopkins University, P50 AG005146, R01 AG020688; Massachusetts General Hospital, P50 AG005134; Mayo Clinic, P50 AG016574; Mount Sinai School of Medicine, P50 AG005138, P01 AG002219; New York University, P30 AG08051, MO1RR00096, UL1 RR029893, 5R01AG012101, 5R01AG022374, 5R01AG013616, 1RC2AG036502, 1R01AG035137; Northwestern University, P30 AG013854; Oregon Health & Science University, P30 AG008017, R01 AG026916; Rush University, P30 AG010161, R01 AG019085, R01 AG15819, R01 AG17917, R01 AG30146; TGen, R01 NS059873; University of Alabama at Birmingham, P50 AG016582, UL1RR02777; University of Arizona, R01 AG031581; University of California, Davis, P30 AG010129; University of California, Irvine, P50 AG016573, P50, P50 AG016575, P50 AG016576, P50 AG016577; University of California, Los Angeles, P50 AG016570; University of California, San Diego, P50 AG005131; University of California, San Francisco, P50 AG023501, P01 AG019724; University of Kentucky, P30 AG028383, AG05144; University of Michigan, P50 AG008671; University of Pennsylvania, P30 AG010124; University of Pittsburgh, P50 AG005133, AG030653; University of Southern California, P50 AG005142; University of Texas Southwestern, P30 AG012300; University of Miami, R01 AG027944, AG010491, AG027944, AG021547, AG019757; University of Washington, P50 AG005136; Vanderbilt University, R01 AG019085; and Washington University, P50 AG005681, P01 AG03991. The Kathleen Price Bryan Brain Bank at Duke University Medical Center is funded by NINDS grant # NS39764, NIMH MH60451 and by Glaxo Smith Kline. Genotyping of the TGEN2 cohort was supported by Kronos Science. The TGen series was also funded by NIA grant AG034504 to AJM, The Banner Alzheimer’s Foundation, The Johnnie B. Byrd Sr. Alzheimer’s Institute, the Medical Research Council, and the state of Arizona and also includes samples from the following sites: Newcastle Brain Tissue Resource (funding via the Medical Research Council, local NHS trusts and Newcastle University), MRC London Brain Bank for Neurodegenerative Diseases (funding via the Medical Research Council),South West Dementia Brain Bank (funding via numerous sources including the Higher Education Funding Council for England (HEFCE), Alzheimer’s Research Trust (ART), BRACE as well as North Bristol NHS Trust Research and Innovation Department and DeNDRoN), The Netherlands Brain Bank (funding via numerous sources including Stichting MS Research, Brain Net Europe, Hersenstichting Nederland Breinbrekend Werk, International Parkinson Fonds, Internationale Stiching Alzheimer Onderzoek), Institut de Neuropatologia, Servei Anatomia Patologica, Universitat de Barcelona. ADNI Funding for ADNI is through the Northern California Institute for Research and Education by grants from Abbott, AstraZeneca AB, Bayer Schering Pharma AG, Bristol-Myers Squibb, Eisai Global Clinical Development, Elan Corporation, Genentech, GE Healthcare, GlaxoSmithKline, Innogenetics, Johnson and Johnson, Eli Lilly and Co., Medpace, Inc., Merck and Co., Inc., Novartis AG, Pfizer Inc, F. Hoffman-La Roche, Schering-Plough, Synarc, Inc., Alzheimer’s Association, Alzheimer’s Drug Discovery Foundation, the Dana Foundation, and by the National Institute of Biomedical Imaging and Bioengineering and NIA grants U01 AG024904, RC2 AG036535, K01 AG030514. We thank Drs. D. Stephen Snyder and Marilyn Miller from NIA who are *ex-officio* ADGC members. Support was also from the Alzheimer’s Association (LAF, IIRG-08-89720; MP-V, IIRG-05-14147) and the US Department of Veterans Affairs Administration, Office of Research and Development, Biomedical Laboratory Research Program. P.S.G.-H. is supported by Wellcome Trust, Howard Hughes Medical Institute, and the Canadian Institute of Health Research. Data for this study were prepared, archived, and distributed by the National Institute on Aging Alzheimer’s Disease Data Storage Site (NIAGADS) at the University of Pennsylvania (U24-AG041689-01), funded by the National Institute on Aging.

---

**Supplementary Table 1.** Differences in rates of cognitive decline and clinical progression for low and high polygenic hazard score (PHS) individuals who tested positive or negative for amyloid or total tau.

|  | Executive function |  | Memory |  | CDR-SB |  |
| --- | --- | --- | --- | --- | --- | --- |
| | $\beta$ (SE) | *p*-value | $\beta$ (SE) | *p*-value | $\beta$ (SE) | *p*-value |
| High PHS/amyloid + | **-0.43 (0.04)** | **<2×10<sup>-16</sup>** | **-0.37 (0.03)** | **<2×10<sup>-16</sup>** | **1.72 (0.12)** | **<2×10<sup>-16</sup>** |
| High PHS/amyloid - | -0.06 (0.04) | 0.17 | **-0.13 (0.03)** | **6.45×10<sup>-4</sup>** | 0.13 (0.14) | 0.37 |
| Low PHS/amyloid + | **-0.27 (0.03)** | **<2×10<sup>-16</sup>** | **-0.28 (0.03)** | **<2×10<sup>-16</sup>** | **0.90 (0.10)** | **<2×10<sup>-16</sup>** |
| Low PHS/amyloid - | **-0.17 (0.03)** | **2.09×10<sup>-11</sup>** | **-0.14 (0.02)** | **1.09×10<sup>-9</sup>** | **0.42 (0.09)** | **1.99×10<sup>-6</sup>** |
| High PHS/total tau + | **-0.46 (0.04)** | **<2×10<sup>-16</sup>** | **-0.38 (0.03)** | **<2×10<sup>-16</sup>** | **1.86 (0.13)** | **<2×10<sup>-16</sup>** |
| High PHS/total tau - | **-0.20 (0.04)** | **2.04×10<sup>-7</sup>** | **-0.19 (0.03)** | **3.36×10<sup>-8</sup>** | **0.69 (0.13)** | **1.34×10<sup>-7</sup>** |
| Low PHS/total tau + | **-0.22 (0.03)** | **6.86×10<sup>-12</sup>** | **-0.22 (0.03)** | **9.33×10<sup>-15</sup>** | **0.70 (0.11)** | **6.45×10<sup>-11</sup>** |
| Low PHS/total tau - | **-0.16 (0.03)** | **1.30×10<sup>-9</sup>** | **-0.13 (0.02)** | **8.49×10<sup>-8</sup>** | **0.32 (0.10)** | **5.33×10<sup>-4</sup>** |

*CDR-SB: Cognitive dementia rating – Sum of boxes.* Significant effects are in bold

